# LC8 enhances 53BP1 foci through heterogeneous bridging of 53BP1 oligomers

**DOI:** 10.1101/2024.09.27.615446

**Authors:** Jesse Howe, Douglas Walker, Kyle Tengler, Maya Sonpatki, Patrick Reardon, Justin W.C. Leung, Elisar J. Barbar

## Abstract

53BP1 is a key player in DNA repair and together with BRCA1 regulate selection of DNA double strand break repair mechanisms. Localization of DNA repair factors to sites of DNA damage by 53BP1 is controlled by its oligomerization domain (OD) and binding to LC8, a hub protein that functions to dimerize >100 clients. Here we show that 53BP1 OD is a trimer, an unusual finding for LC8 clients which are all dimers or tetramers. As a trimer, 53BP1 forms a heterogeneous mixture of complexes when bound to dimeric LC8 with the largest mass corresponding to a dimer-of-trimers bridged by 3 LC8 dimers. Analytical ultracentrifugation and isothermal titration calorimetry demonstrate that only the second of the three LC8 recognition motifs is necessary for a stable bridged complex. The stability of the bridged complex is tuned by multivalency, binding specificity of the second LC8 site, and the length of the linker separating the LC8 binding domain and OD. 53BP1 mutants deficient in bridged species fail to impact 53BP1 focus formation in human cell culture studies, suggesting that the primary role of LC8 is to bridge 53BP1 trimers which in turn promotes recruitment of 53BP1 at sites of DNA damage. We propose that the formation of higher-order oligomers of 53BP1 explains how LC8 elicits an improvement in 53BP1 foci and affects the structure and functions of 53BP1.

## Introduction

Anti-tumor protein p53 binding protein (53BP1) is a 200 kDa protein involved in DNA repair and cell cycle regulation^1–3^. Together, breast cancer type 1 susceptibility gene (BRCA1) and 53BP1 regulate pathway selection for DNA double strand break repair^4^. 53BP1 binds and protects DNA ends formed by double strand breaks, limiting end resection and promoting use of nonhomologous end joining (NHEJ) for DNA repair^2,5,6^. BRCA1 is necessary for eviction of 53BP1 from chromatin, allowing for end resection factors to initiate homology directed repair (HDR)^7^. In the absence of functional BRCA1, the HDR pathway is nonfunctional and error prone NHEJ dominates double strand break repair^8^. Aberrant NHEJ results in genome instability such as chromosomal rearrangements and radial chromosome formation. Chemotherapy drugs such as PARPi and platinum-based drugs are effective against BRCA negative cancers^2,6,9,10^. However, in BRCA negative cells, inactivation of 53BP1 once again activates HDR, resulting in loss of efficacy of PARPi and other chemotherapy agents. Understanding the functional states of 53BP1 is therefore critical to creating effective drugs and treatment strategies for cancer patients.

In response to DNA damage, 53BP1 forms nuclear bodies which mark sites of DNA double strand breaks^6,11,12^. Upon induction of DNA damage, 53BP1 is recruited to sites of DNA double strand break and is retained on chromatin through interactions between modified histones and the tudor domain^13^, ubiquitin dependent recognition domain^14^ and oligomerization domain^15^ (Fig. 1A). The long N-terminal disordered domain of 53BP1 (residues 1-1230), which contains 28 putative ATM kinase recognition (S/TQ) motifs, is phosphorylated by ATM kinase at multiple sites ^16,17^, which results in recruitment of downstream effectors active in protection of DNA ends (e.g. RIF1^18^, Rev7^19^ and Shieldin^20^) or in NHEJ (e.g. PTIP^21^).The recruitment of 53BP1 to nuclear bodies is critical to its function in DNA repair.

**Figure 1.**
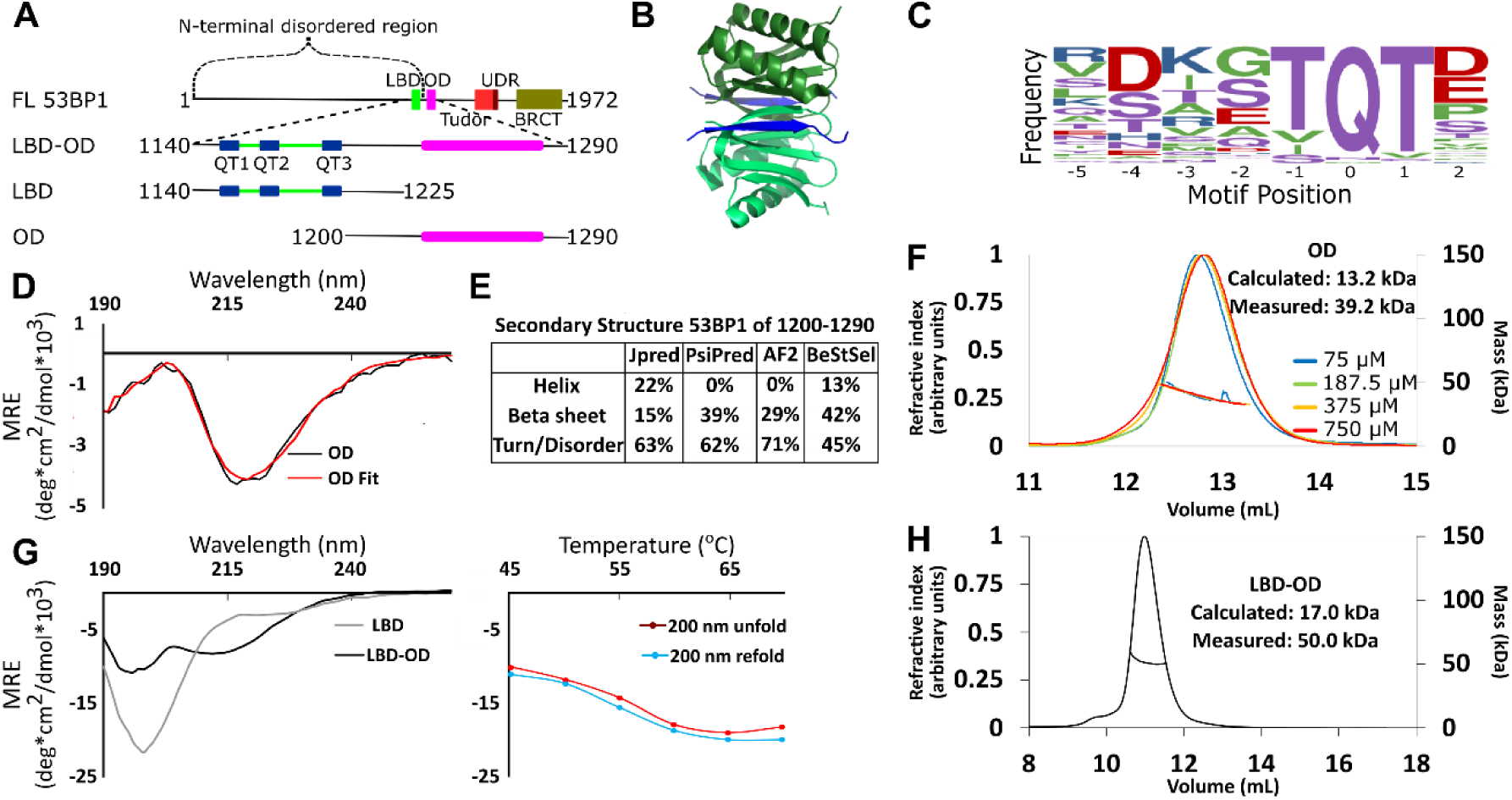
Domain architecture of 53BP1 and preliminary characterization. **A**, Map of 53BP1 domain architecture showing LC8 binding sites (QT1-3) in blue and the OD in magenta. **B**, Structure of LC8 dimer (green) bound to two strands a peptide partner (blue) (PDB code 5E0L). **C**, Variable binding motif recognized by LC8. The x-axis shows the residue position and y-axis shows the frequency of residues found in known LC8 binding clients, 111 used for this sequence logo. The anchor is the least variable, and is assigned residue numbers of -1, 0, and 1. **D**, Circular dichroism data for 53BP1 OD (aa 1200-1290) overlayed by BeStSel fit. Data are presented in units of mean residue ellipticity (MRE). Percent secondary structure is reported in *F*. **E**, Secondary structure predictions of 53BP1 1220-1290 using JPred, PsiPred, and AlphaFold2. **F**, Percentage of secondary structure of 53BP1 OD determined from JPred, PsiPred, AlpaFold2, and BesStSel fit of CD data. **G**, (Left) Circular dichroism of 53BP1 LBD-OD overlaid with LBD spectra, shown here for comparison. In addition to the minima at 200 nm for LBD-OD, indicative of disorder, the minima near 218 nm shows evidence of an ordered region. (Right), thermal denaturation (red) and refolding (blue) of LBD-OD showing increase in ellipticity at 200 nm. **H**, SEC-MALS of 53BP1 LBD-OD shows mass suggesting a trimer.

53BP1 contains 3 dynein light chain 8 (LC8) binding sites at the C-terminus of its disordered domain (residues 1150-1230)^22,23^, followed by the oligomerization domain^13^ (residues 1231-1272) which is poorly characterized. LC8 is a hub protein that binds >100 client proteins at their disordered regions in its two symmetric binding grooves, facilitating the dimerization of the client proteins^24–26^ (Fig. 1B). Interplay between the oligomerization domain (OD) and LC8 binding domain (LBD) is crucial to function of 53BP1^6,10^. Either the OD mutant (ODm) or LC8 binding mutant (LC8m) reduces 53BP1 nuclear bodies and a construct containing both mutations reduces nuclear bodies to levels similar to the control^6^. Furthermore, the LC8m shows reduced DNA repair activity, while the ODm is inactive in DNA repair and the LC8m/ODm construct is not even efficiently recruited to nuclear bodies. The functional synergy between the OD and LC8 binding in formation of 53BP1 nuclear bodies supports a model in which LC8 either stabilizes dimeric 53BP1 to increase affinity for bivalent interaction with chromatin as proposed^6^ or that LC8 stimulates higher order oligomerization in 53BP1^27^.

LC8 binds a highly variable short linear recognition motif in disordered regions of client proteins (Fig. 1C)^24,25^. A well conserved triad, TQT, most frequently occupies the anchor of this motif, but the entire sequence is variable. Due to the strong preference for binding to QT motifs, we refer to LC8 binding sites as QT sites (or QTs). Both the oligomerization domain of 53BP1 and the binding of the three QTs to dimeric LC8 are expected to contribute to oligomerization of 53BP1. Intriguing about our work here is the discovery that 53BP1 is a trimeric LC8 client (Fig. 1H), leading to the question of how dimeric LC8 binds a trimeric client and the functional role of this binding. Well-described mechanisms through which LC8 regulates oligomeric clients include regulation of the affinity of self-association^28,29^, structuring of the disordered region^30,31^, and alignment of domains by restriction of the conformational ensemble of the client^32^. Here, we show a novel binding mode for LC8, bridging of trimeric client subcomplexes, and propose that LC8 improves 53BP1 focus formation through heterogeneous bridging of 53BP1 subcomplexes.

## Results

### Structural Characterization of LBD-OD

The oligomerization domain (OD) of 53BP1 spans residues 1231-1279 based on truncation studies^13^. Disorder prediction by IUPred^33^ suggests that residues 1240-1270 are ordered. Since coiled-coils are the most common type of oligomerization domain in LC8 binding clients^29,34^, we ran predictions for coiled-coil propensity. Neither Waggawagga^35^ nor Marcoil^36^ predict any coiled-coils in this region. Circular dichroism (CD) of a construct of the OD spanning residues 1200-1290 shows a single minimum near 218 nm (Fig. 1D) consistent with a primarily beta-sheet structure. Fitting this data with BeStSel^37^ estimates a primarily beta-sheet structure with some propensity for an alpha helical structure (Fig. 1E). Structure prediction algorithms are in general agreement in the structured regions (Fig. 1E). AlphaFold2^38^ and PsiPred^39^ predict an entirely beta-sheet structure while JPred^40^ predicts a mixture of alpha-helices and beta-strands. Since the BeStSel CD-based structure analysis estimates a higher percentage of structure than the sequence-based predictions, we conclude that the increased structure is due to some folding upon oligomerization (Fig. 1E).

The LBD remains disordered in the context of the OD as shown by a sharp peak near 200 nm and a smaller minimum around 218 nm in the CD spectrum consistent with a primarily disordered structure containing some beta-sheet structure (Fig. 1G). The spectrum of LBD alone (residues 1140-1225) contains only the minimum at 200 nm, indicating the ordered domain of LBD-OD is between residues 1225 and 1290 (Fig. 1G)^13^. Thermal denaturation assays of LBD-OD by CD show a decrease in the intensity of the minimum near 215 nm and a simultaneous increase in the minimum at 200 nm (Fig. 1G). Both LBD-OD (Fig. 1G) and OD (data not shown) have a T_m_ at 57 °C. Refolding curves show no hysteresis indicating that both LBD-OD and OD refold reversibly.

Size-exclusion chromatography coupled to multi-angle light scattering (SEC-MALS) of the OD (1200-1290) shows a mass of 39.2±0.6 kDa as an average of 4 runs at concentrations between 75-750 μM (Fig. 1F), which suggests a trimeric form dominates in this concentration range (expected trimer mass of 39.5 kDa) LBD-OD shows a single peak with mass of 50 kDa (Fig. 1H). Since the calculated mass expected for a monomeric LBD-OD is 17.0 kDa, the mass measured by SEC-MALS suggests that 53BP1 OD is a trimer (expected mass of 51 kDa for trimer). To date, all known oligomeric clients of LC8 are either dimers or tetramers, and thus the mechanism for complex formation between a dimer and a trimer is not known.

### Disorder of the LBD is retained in the LBD-OD

The ^1^H-^15^N HSQC of LBD-OD shows low peak dispersion and overlays well with the HSQC of LBD (Fig. 2A). No additional peaks for the OD were detected in the concentration range of 50 to 400 μM and in the temperature range of 10-40 °C. Cα-Cβ chemical shift indexing of LBD and LBD-OD shows shifts less than 0.5 for most residues, indicating that both constructs are disordered between residues 1140-1225 (data not shown). Most residues of the LBD in the context of the OD show similar dynamic behavior to that of the LBD alone, indicating that the LBD remains disordered in the LBD-OD construct (Fig. 2B-E). However, residues 1210-1225, which are at the end of the LBD, appear less dynamic with high R2 values (Fig. 2C) compared to those for the same residues in the LBD alone as expected due to its proximity to a folded domain. Taken together, these data indicate that the LBD-OD retains flexibility in the intrinsically disordered LBD but possesses a folded region that corresponds to the OD.

**Figure 2.**
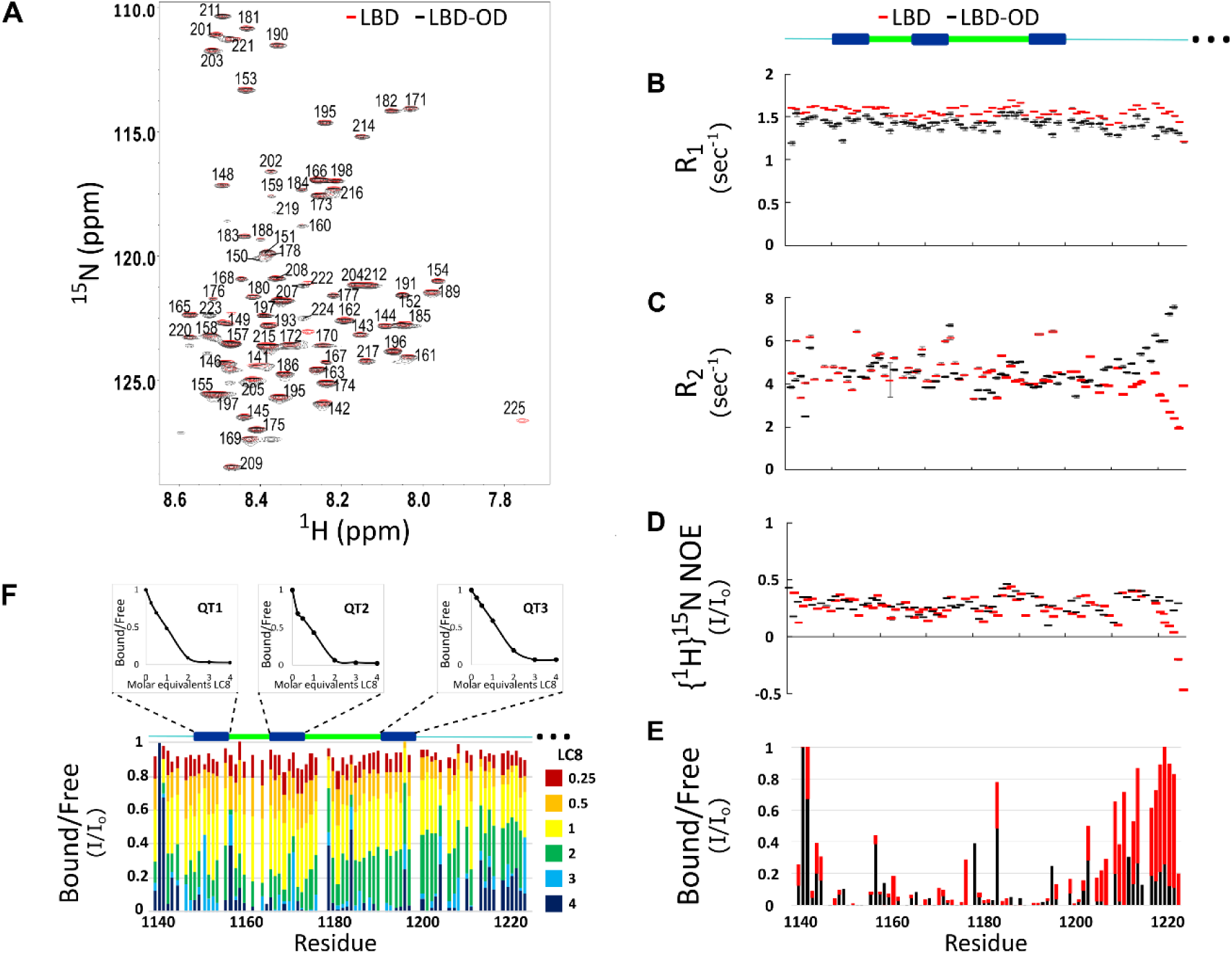
Comparison of structure and interactions of LBD and LBD-OD with LC8 probed by NMR. **A**, Overlay of ^1^H-^15^N HSQC spectra of 53BP1 LBD (red) and LBD-OD (black) acquired at 800 MHz at 10°C in 20 μM sodium phosphate (pH 6.5) and 50 μM sodium chloride with 10% D_2_O. The residues are labeled without the thousands place, which is 1--- for every residue to minimize crowding. Resonances overlay well for almost all residues. **B**, R1, **C,** R2, and **D** {^1^H}^15^N NOE overlay of LBD (red) and LBD-OD (black). Diagram of 53BP1 LBD with LC8 QT sites labeled in blue shown at the top. **E,** Overlay of ^15^N LBD and ^15^N LBD-OD peak height in ^1^H-^15^N HSQC in the presence of 4 molar equivalent of LC8. **F**, Titration of uniformly labeled ^15^N LBD-OD with unlabeled LC8. A diagram of LBD-OD is shown above the plot which highlight the locations of QT sites (blue). The average signal for the 8 residues comprising the QT site is shown in decay curves above each QT site as a function of LC8 concentration.

Titration of LBD with LC8 shows uniform signal attenuation in residues 1150-1200 of 53BP1, suggesting no binding preference for any particular site^23^. Titration of LBD-OD with LC8 shows similar uniform signal attenuation for the same residues (Fig. 2F) suggesting that the cooperative binding of LBD is conserved in the presence of the OD.

### 53BP1 OD forms heterogeneous complexes when bound to LC8

The interaction between LBD and LC8 by analytical SEC and NMR support a model in which LBD binds LC8 cooperatively in a single step^23^. SEC is shown here for comparison (Fig. 3A). In contrast, SEC-MALS of LBD-OD with excess LC8 shows a heterogenous complex containing at least three major populations (Fig. 3B) with masses in the 120-296 kDa range (Table 1). Dimeric LBD-OD with 3 LC8 dimers has a calculated mass of 95 kDa, much smaller than what we observed. A complex containing one LBD-OD trimer and 3 LC8 dimers is expected to have a mass of 114 kDa, and a complex containing 2 LBD-OD trimers and 9 LC8 dimers would have mass of 293 KDa (Table 2) which are both consistent with the measured masses. Potential intermediates that contain 2 LBD-OD trimers could occupy between 3 and 9 dimers of LC8. Our SEC-MALS suggests that LBD-OD forms a trimeric complex with 3 dimers of LC8 sites and a bridged complex containing 2 trimers of LBD-OD and up to 9 dimers of LC8 (Fig. 3D). For comparison, the LBD alone which shows a much simpler titration behavior (Fig. 3A) with a single peak for the LBD-LC8 complex implying a single step to form a duplex complex (Fig. 3C).

**Figure 3.**
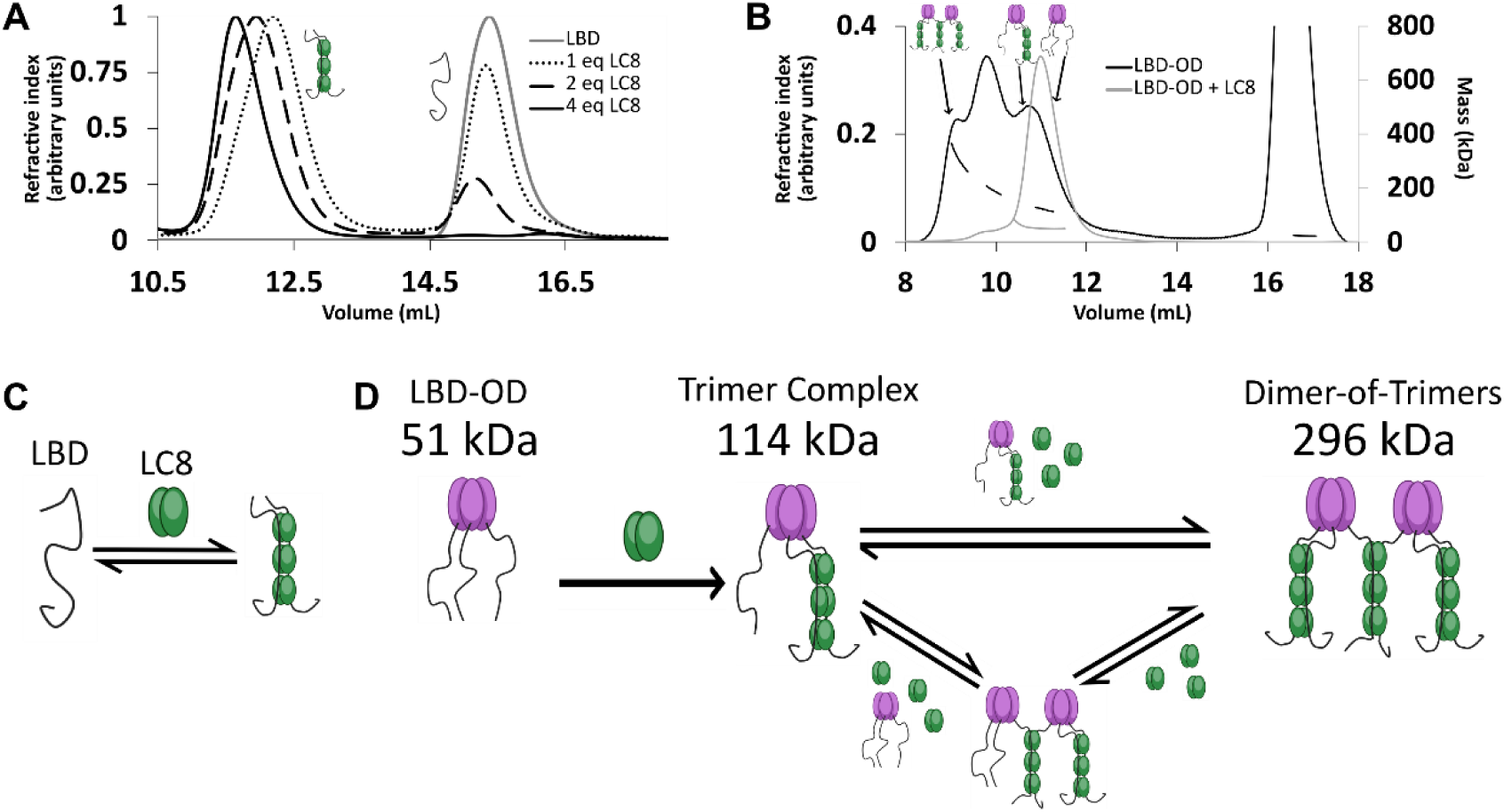
LBD-OD forms a heterogeneous complex when bound to LC8. **A**, Titration of 53BP1 LBD results in a single species with large mass, even at low stoichiometric equivalents. Data is adapted from Howe et. al. 2022^23^ and shown here for comparison. **B**, SEC-MALS of free LBD-OD and LBD-OD with 4 molar equivalents of LC8 show formation of a heterogeneous mixture of complexes with masses in the 120-296 kDa range (Table 2). **C**, Model of LBD binding LC8. A duplex containing 2 strands of LBD and 3 dimers of LC8 is created in a single step. **D**, Model of LBD-OD binding LC8. 53BP1 trimers bind LC8 forming a trimer complex containing one trimer of 53BP1 and 3 LC8 dimers. The trimer complex binds excess LC8 forming bridged complexes.

**Table 1.** Masses measured by SEC-MALS.

| Measured |  |  |  |
| --- | --- | --- | --- |
| Construct | High mass (kDa) | Intermediate mass (kDa) | Low mass (kDa) |
| LBD-OD | 296±2.5 | 190±1.7 | 120±2.2 |
| QT2-OD | 168±1.2 | 104±0.6 | 80±.5 |
| QT3-OD | - | - | 69±0.4 |
| QT12-OD | 247±1.2 | 178±0.8 | 112±0.5 |
| QT13-OD | - | - | 92±0.5 |
| QT23-OD | 233±1.3 | 123±0.7 | 101±0.6 |

**Table 2.** Expected masses of LBD-OD mutants in different LC8 complexes.

| Complex | Bridged complex ratio<br>(53BP1 trimers:<br>LC8 dimers) | Expected mass<br>(kDa) | Trimer complex<br>(53BP1 trimers:<br>LC8 dimers) | Expected mass<br>(kDa) |
| --- | --- | --- | --- | --- |
| LBD-OD | 2:9 | 292.8 | 1:3 | 114.6 |
| One-site mutants | 2:3 | 165.6 | 1:1 | 72.2 |
| Two-site mutants | 2:6 | 229.2 | 1:2 | 93.4 |

### Structural basis of heterogeneity in LBD-OD complexes with LC8

To determine the importance of multivalency in 53BP1-LC8 interactions, we generated a series of mutants which change the three anchor residues of the LC8 binding motif to alanine (AAA), eliminating LC8 binding at the sites of the mutations. We name each construct by the sites left intact (e.g. QT13-OD has AAA mutation in QT2, leaving QTs 1 and 3 to bind LC8) (Fig. 4A).

**Figure 4.**
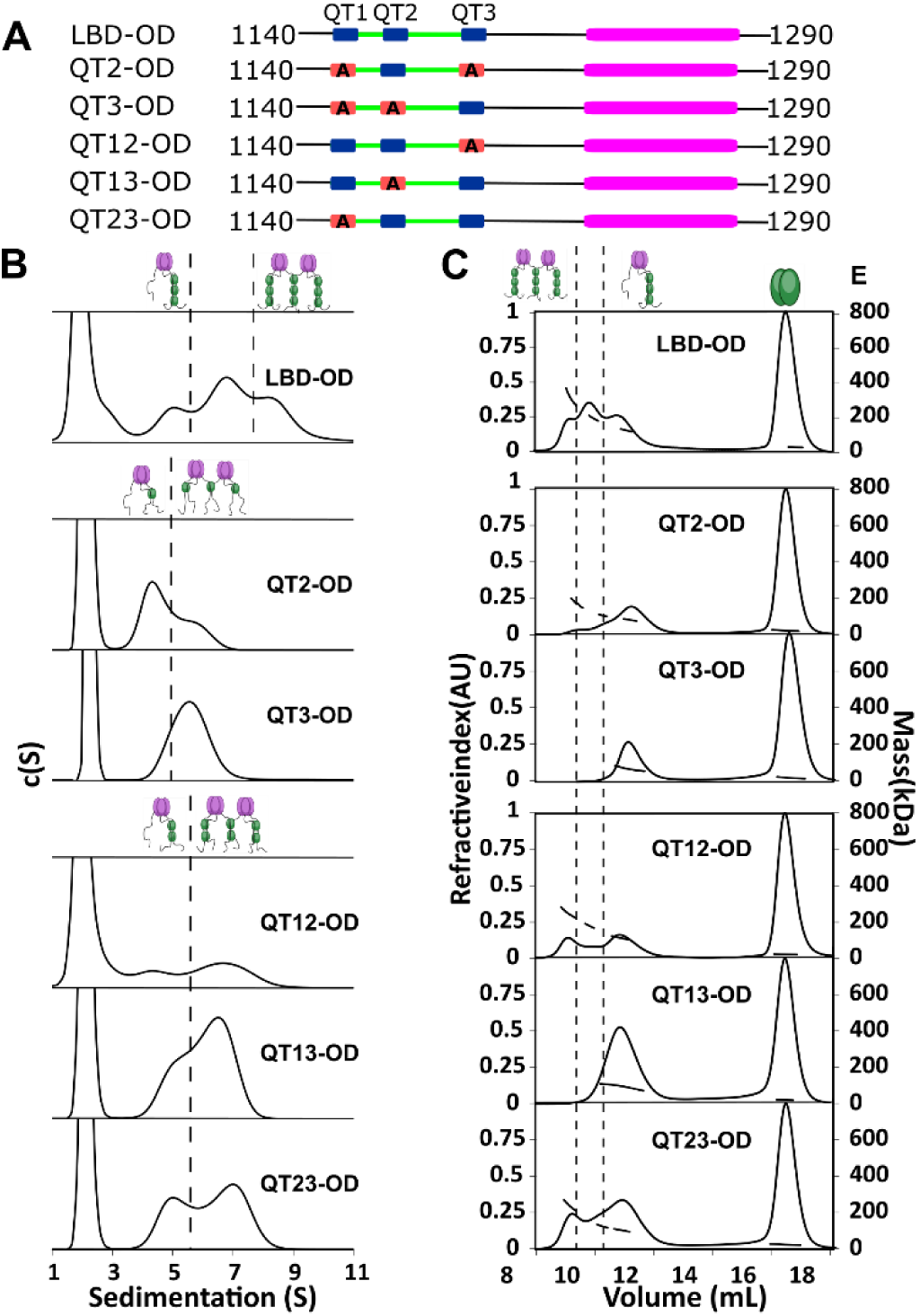
QT2 determines stability of bridged complex. **A**, Domain maps of LBD-OD QT mutants. LC8 binding sites (QT) are shown in blue, while QTs mutated to abolish binding are shown in red. **B**, SV-AUC of WT (top) one-site mutants (center) and two-site mutants (bottom) in 50 mM sodium phosphate and 150 mM sodium chloride at 25°C. Dotted lines separate large mass peaks from small mass peaks. Peaks with high S value are assigned to the dimer-of-trimers complex and low mass peaks to the trimer complex, represented in cartoon. **C**, SEC-MALS of complexes formed by LBD-OD and mutants in 50 mM sodium phosphate and 150 mM sodium chloride (pH 7.2). Dotted lines separate large mass peaks from small mass peaks. In SEC-MALS experiments, dimer-of-trimers complexes are only observed for the complexes containing intact QT2.

In SV-AUC of one-site LBD-OD mutants (Fig. 4B), two peaks were observed, presumably corresponding to trimeric and bridged complexes. QT3-OD primarily formed bridged complexes, while QT2-OD exhibited a mixture of trimeric and bridged species. In two-site QT mutants (Fig. 4B), those containing QT3 exhibited a higher proportion of bridged complexes, with QT13-OD showing the highest proportion of complex with high S value.

SEC-MALS analysis identified the mutants that form the most stable bridged complexes with LC8 that survived dilution on the SEC column (Fig. 4C). In contrast to SV-AUC, QT3-OD is exclusively in the trimeric state, while QT2-OD is mostly trimeric but retains a small fraction of bridged complexes. QT1-OD dissociated nearly to completion, and so we do not include it in our analysis. This difference between SEC and SV-AUC profiles suggests that while both QT2 and QT3 mutants can form bridged complexes with excess LC8 as observed by SV-AUC, only QT2-OD maintains stable bridging upon dilution during SEC. Comparison of expected and measured masses further demonstrated that complexes containing intact QT2 have populations with masses consistent with the bridged complex. These findings suggest that QT2 is necessary to form stable intermolecular interactions resulting in intact bridged complexes.

### Thermodynamics analysis of LBD-OD:LC8 interaction reveal a unique role for QT2

Isothermal titration calorimetry (ITC) showed different binding affinities for each site in the LBD^23^. Importantly, QT2 is the only site with an unfavorable entropy and QT1 is the weakest binding site overall. Here, we investigate LC8 binding to 53BP1 in context of the OD.

Analysis of isotherms from LC8 titrated into LBD-OD and the one- and two-site mutants with the one-set-of-sites (OSS) model shows enthalpically favorable interactions with K_d_ in the sub-micromolar to low micromolar range (Fig. 5K, Table 3). The wild-type LBD-OD (Fig. 5A) shows a stoichiometry near 3:1 (3 LC8: 1 53BP1; or 9 LC8 dimers and two trimers of 53BP1), suggesting that a bridged complex could be formed and that all the LC8 sites are filled.

**Figure 5.**
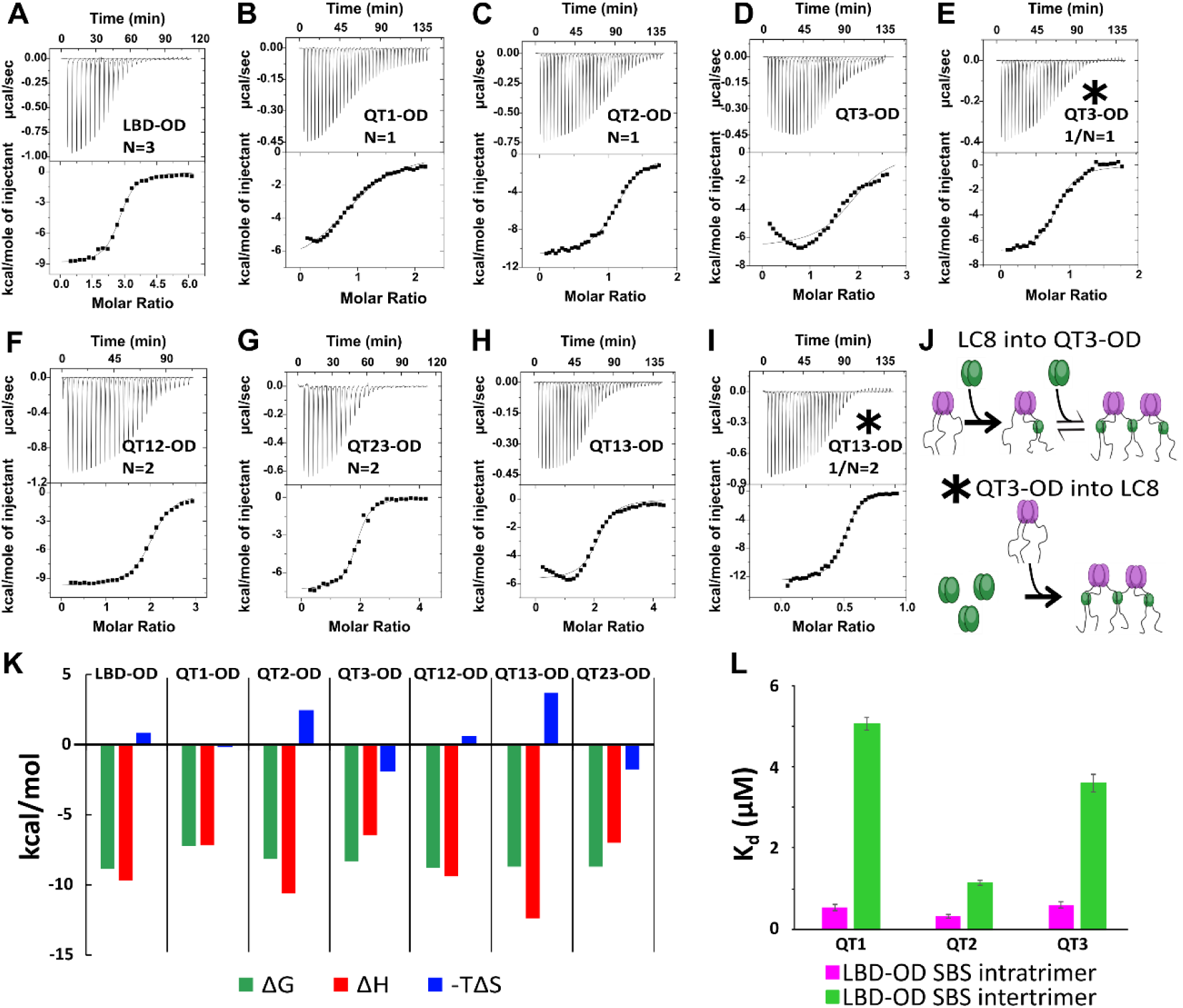
Thermodynamics of interactions of LBD-OD and mutants with LC8. **A-D, F-H**, Representative isotherms for LBD-OD (A), QT1-OD (B), QT2-OD (C), QT3-OD (D), QT12-OD (F), QT13-OD (G), QT23-OD (H) with LC8 in 50 mM sodium phosphate and 150 mM sodium chloride (pH 7.2) at 25°C. In these experiments, LC8 at 250-400 μM was titrated into LBD-OD or mutant at 10-30 μM. **E,I**, Isotherm of QT3-OD (E) and QT13-OD (I) at 200 μM titrated into 20 μM LC8. The downward inflection early in the titration is not present in this isotherm, as it is in the titration of LC8 into mutant in D and H. **J**, A cartoon illustrating how a stable intermediate is formed during titration. (Top) When LC8 is titrated into QT3-OD, excess 53BP1 binds LC8 resulting in forming the trimeric intermediate. (Bottom) When QT3-OD is titrated into LC8, excess LC8 allows the bridged complex to form without a stable intermediate. **K**, Bar graph of thermodynamic parameters for A-G isotherms fit to OSS model. Thermodynamic parameters are shown in Table **L**, Affinities of one-site binding in 53BP1 determined from fits to the OSS model for LBD and LBD-OD are in gray and orange and using subsequent binding sites (SBS) model in cyan (intratrimer) and magenta (bridging). QT2-OD has the smallest difference in affinity between intratrimer and bridging interactions.

**Table 3.**
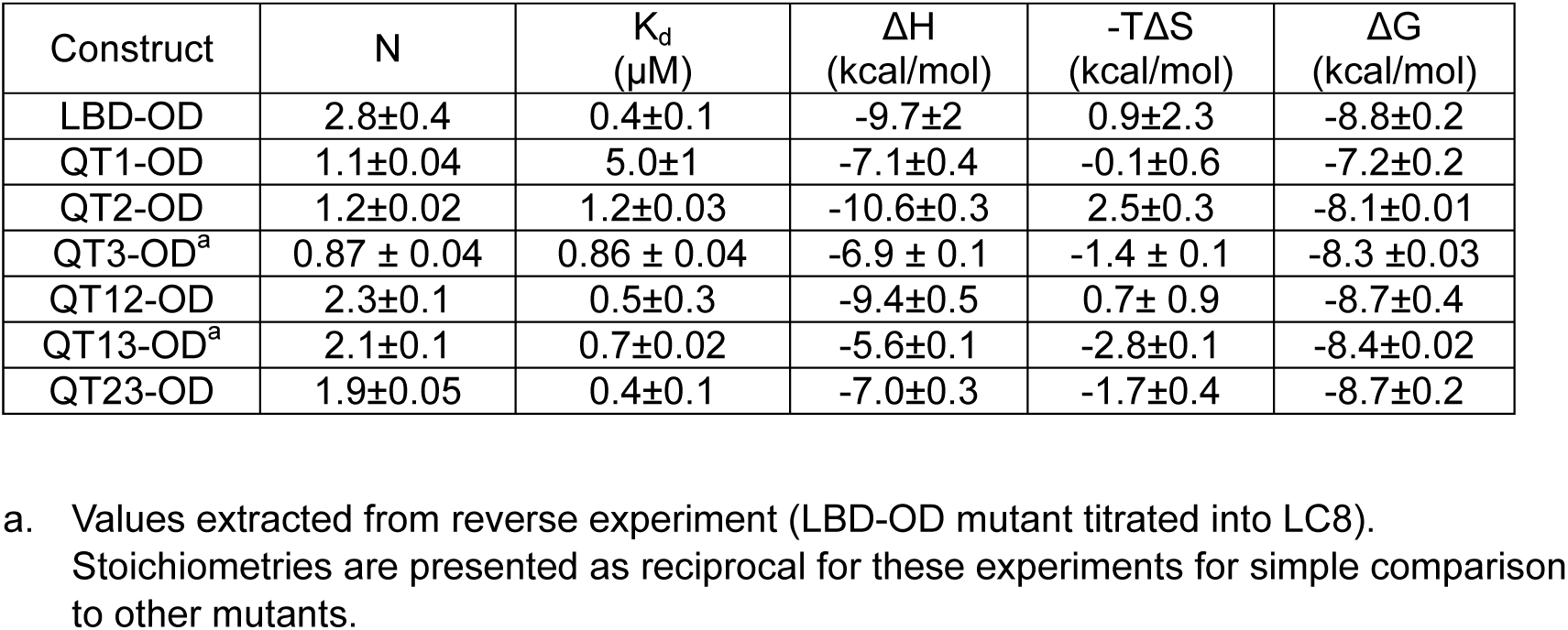
Thermodynamics of LBD-OD:LC8 interactions measured by ITC.

| Construct | N | K<sub>d</sub><br>(μM) | ΔH<br>(kcal/mol) | -TΔS<br>(kcal/mol) | ΔG<br>(kcal/mol) |
| --- | --- | --- | --- | --- | --- |
| LBD-OD | 2.8±0.4 | 0.4±0.1 | -9.7±2 | 0.9±2.3 | -8.8±0.2 |
| QT1-OD | 1.1±0.04 | 5.0±1 | -7.1±0.4 | -0.1±0.6 | -7.2±0.2 |
| QT2-OD | 1.2±0.02 | 1.2±0.03 | -10.6±0.3 | 2.5±0.3 | -8.1±0.01 |
| QT3-OD<sup>a</sup> | 0.87 ± 0.04 | 0.86 ± 0.04 | -6.9 ± 0.1 | -1.4 ± 0.1 | -8.3 ± 0.03 |
| QT12-OD | 2.3±0.1 | 0.5±0.3 | -9.4±0.5 | 0.7± 0.9 | -8.7±0.4 |
| QT13-OD<sup>a</sup> | 2.1±0.1 | 0.7±0.02 | -5.6±0.1 | -2.8±0.1 | -8.4±0.02 |
| QT23-OD | 1.9±0.05 | 0.4±0.1 | -7.0±0.3 | -1.7±0.4 | -8.7±0.2 |

a. Values extracted from reverse experiment (LBD-OD mutant titrated into LC8). Stoichiometries are presented as reciprocal for these experiments for simple comparison to other mutants.

The one-site LBD-OD mutants (Fig. 5B-E) have similar binding properties to LBD^23^ in that QT2 is the only one-site mutant with unfavorable entropy. Interestingly, both QT1 and QT3 binding gives non-sigmoidal isotherms that we attribute to multi-step binding. In these constructs, two of the pre-associated chains of the trimer bind to LC8 first, leaving a third LBD chain available to bridge the available LBD chain from another trimer (Fig. 5J). To test this model, we reversed the direction of titration, resulting in an excess of LC8 in early titration points which we hypothesized would push the reaction such that each QT3-OD binding site was fully bound at every injection. Titrating QT3-OD into LC8 produced a perfectly sigmoidal isotherm indicating that no intermediate is formed in line with our model (Fig. 5E).

The resulting thermodynamic parameters as calculated by the OSS model are based on a convolution of the stronger intratrimeric binding and the weaker intertrimeric (bridging) LC8 binding and therefore is a weighted average of the two events. The absence of a non-sigmoidal isotherm when titrating LC8 into QT2-OD can be explained as being due to the intermediate being only briefly occupied, suggesting that the bridging interaction has a stronger binding affinity closer to that of the intratrimeric LC8 binding than that of the other single site mutants. We modelled our system as a hexamer with three binding events (two intratrimer events and one bridging event) using a subsequent binding sites (SBS) model in Origin 7.0 (Fig. 5L), which enables extracting thermodynamics of multiple binding events. Using this model, we identify two clearly different binding affinities in the one-site LBD-OD mutants. We assign the lower affinity binding event to the bridging of 53BP1 trimers and the higher affinity to the intratrimer interactions. For the intratrimer interactions, the difference in affinity for each mutant is very small (K_d_ between 0.3 and 0.6 µM), while the difference between affinities for bridging interactions is larger (K_d_ between 1 and 5 µM). The mutant with the smallest difference between bridging affinity and intratrimer interactions is QT2-OD (intratrimer K_d_ = 0.3 µM, intertrimer K_d_ = 1.1 µM). This small difference explains the absence of intermediate in ITC isotherms for LBD-OD containing intact QT2. Additionally, the fact that QT2-OD has the strongest affinity for the bridging interaction provides a mechanistic explanation for our observation that QT2 is essential for a strongly bridged dimer-of-trimers complex. Overall, the SBS fitting of LBD-OD one-site mutants isotherms supports a model in which trimers of 53BP1 first bind LC8 as a trimer before forming a bridged species.

To better understand the interplay between multiple LC8 sites in 53BP1, we performed ITC on two-site mutants of LBD-OD (Fig. 5 F-I). All interactions have affinity around 0.5 μM, but QT13-OD, the only two-site mutant not containing QT2, gives a non-sigmoidal isotherm. Again, when we reverse the direction of titration (QT13-OD into LC8), the isotherm becomes sigmoidal (Fig. 5H, I). Since all LBD-OD mutants containing QT2 have sigmoidal isotherms, we attribute the non-sigmoidal isotherms to the formation of a trimeric intermediate and suggest that QT2 binding may pay the entropic penalty for bridging 53BP1 trimers.

### Reducing linker length between LBD and OD stabilizes dimer-of-trimers

For ASCIZ, which binds LC8 multivalently to regulate transcription, linker length contributes to both compositional^41^ and conformational^42^ heterogeneity. To determine a possible similar effect for linker length on 53BP1 heterogeneity, we focused on the linker between 53BP1 LBD and OD, hypothesizing that the LBD-OD linker would regulate the bridging interaction. We generated two mutants of LBD-OD by deleting 5 and 15 residues (Δ1206-1210 and Δ1206-1220, respectively) (Fig. 6A) in the ∼30 residue-long linker. Shortening the linker by 5 residues results in a population shift to the bridged complex and further reducing the length by 15 residues shifts the population to an almost completely bridged complex (Fig. 6B) suggesting that shortening the LBD-OD linker stabilizes the dimer-of-trimers complex (Fig. 6C). In support of this model, ITC shows sigmoidal isotherms for both linker deletion mutants (Fig. 6D). The entropic contribution becomes more favorable as the linker length is decreased while the overall affinity of the interaction is unchanged (Fig. 6E).

**Figure 6.**
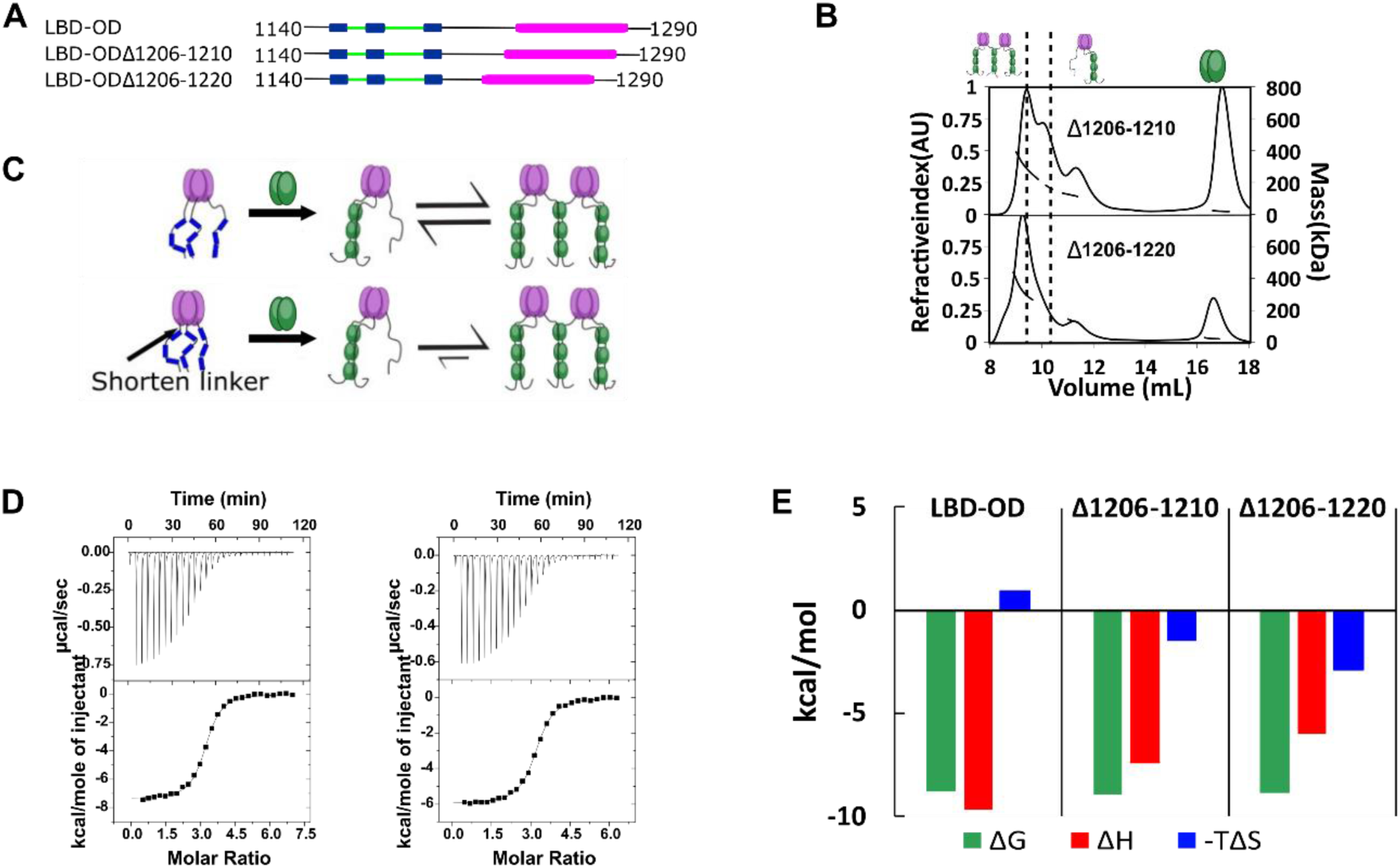
Linker between LBD and OD determines stability of bridged complex. **A**, Domain map of 53BP1 linker deletion mutants. **B**, SEC-MALS of 53BP1 linker deletion mutants show high proportion of bridged complexes. **C**, Model of the effect of reducing the linker in 53BP1 LBD-OD. Shorter linker results in increased population of bridged complex seen by SEC-MALS. **D**, ITC shows sigmoidal curves with stoichiometry near 3, suggesting that the linker deletion mutants form bridged complex in a single binding step. **E**, The entropic contribution becomes more favorable as the linker length is decreased, but the overall affinity of the interaction is unchanged.

### QT2 sequence determines proportion of bridged complex

The effect of sequence specificity in LC8 binding has been studied in great detail, revealing a variable motif which binds LC8 with varying affinities depending on the sequence of the client^24,25,34,43,44^. Most important to the affinity of LC8 binding are three anchor residues within the LC8 binding motif (Fig. 7A). Mutations within the LC8 binding anchor have large impacts on binding, but the -1 position exhibits the most variability (Fig. 1B). Within 53BP1, all three QTs contain a different residue in the -1 position (Fig. 7A, bottom). Two LBD-OD QT2 variants which are still expected to bind LC8 (T1171I and T1171V), and a phosphomimetic variant which is expected to greatly reduce binding for QT2 (T1171E) and alter the populations of LBD-OD complexes observed in SEC-MALS (Fig. 7B). As expected, T1171E elutes completely as a trimer complex, like QT13-OD, while both T117I and T1171V form a mixture of trimer and bridged complexes. T1171I forms similar population of trimer and bridged complex to the WT, while T1171V forms a significantly higher population of bridged complex (Fig. 7B), indicating that sequence specificity of QT2 is a major determinant of 53BP1 heterogeneity. BLAST analysis of the human 53BP1 sequence shows that the anchor of QT2 is well conserved in animals, in support of a critical role for 53BP1-LC8 interactions being tuned to maintain heterogeneity of 53BP1 oligomerization.

**Figure 7.**
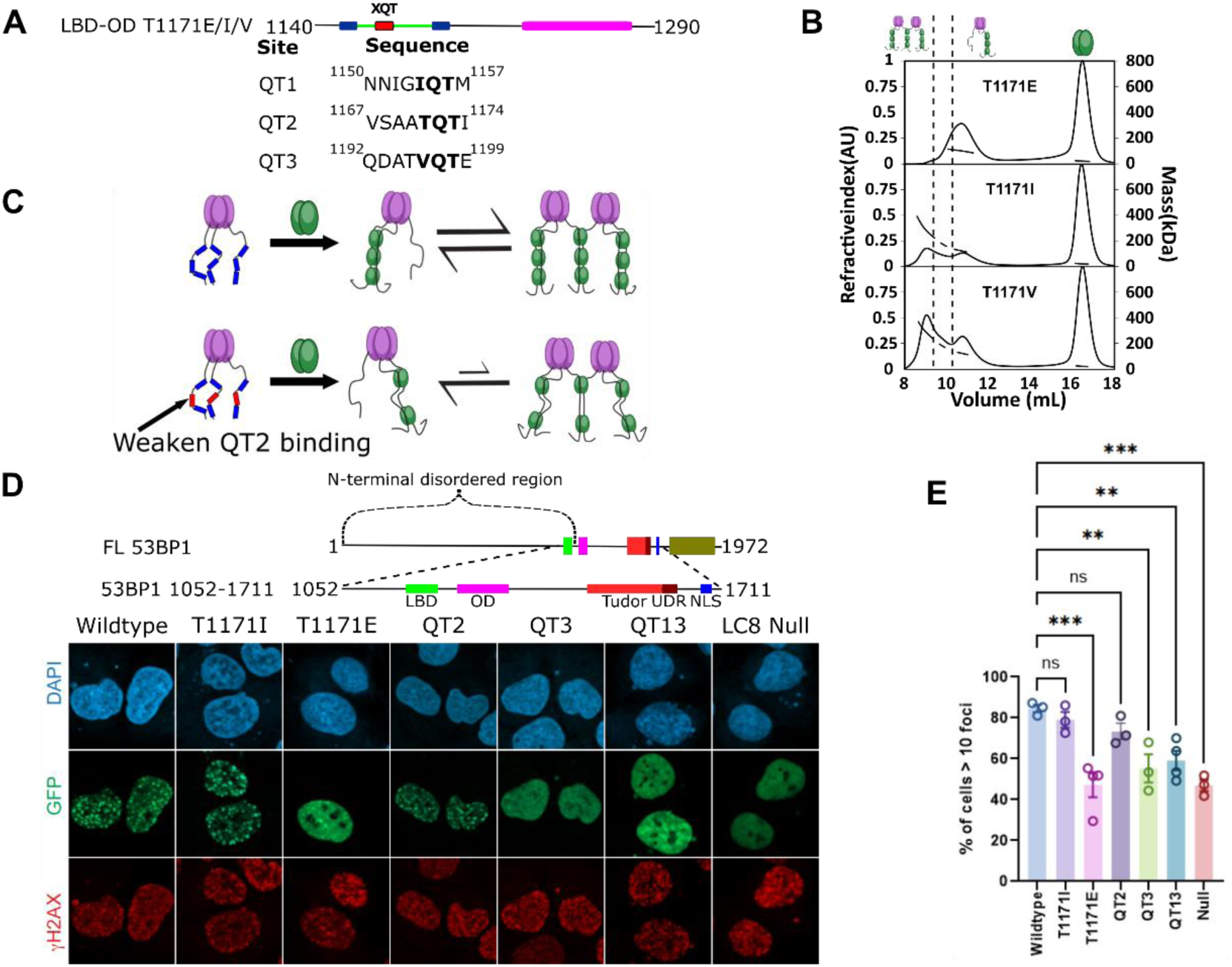
LC8 bridging promotes 53BP1 recruitment to DNA damage sites. **A**, (Top) Domain map of 53BP1 QT2 mutants. (Bottom) Sequence of the three QTs in 53BP1. QT2 is the only site with the canonical TQT anchor. **B**, Mutation of the -1 site (T1171) in 53BP1 has drastic effects on the relative population of trimer and bridged complexes. The phosphomimetic (T1771E) elutes completely as a trimer, while both phosphodeficient mutants (T1171I/V) generate a mixture of trimer and bridged complex. **C**, Model showing the effect of reducing the linker length in 53BP1 LBD-OD on the proportion of bridged complex. **D**, (top) Domain map of constructs transfected into U2OS cells. These constructs span residues 1052-1711 and contain the LBD, OD, Tudor, UDR, and nuclear localizations sequence (NLS) of 53BP1. (bottom) Representative immunofluorescence micrographs of GFP-labeled 53BP1 fragment a.a.1052-1711 wildtype and mutants, at 1 hour after 10 Gy irradiation. **E**. Quantification of 53BP1 foci as presented in D for the indicated expression vectors. The error bars correspond to mean±S.E.M.

### Ability to form a bridged complex is correlated with enhancement in 53BP1 focus formation

53BP1 contains a minimal focus forming (MFF) region spanning residues 1220-1711, which includes the OD, tandem tudor domain, ubiquitin-dependent recruitment domain, and nuclear localization sequence^13^ (Fig. 7D). While LC8 binding is not required for focus formation^12^, it improves focus formation and recruitment of 53BP1 to chromatin^6,10^. A series of 53BP1 mutants spanning the LBD and MFF were tested for their ability to improve 53BP1 focus formation in response to ionizing radiation. The phosphodeficient mutant (T1171I) produces an enhancement in 53BP1 foci similar to the wildtype protein (79±7% for T1171I vs 85±3% for wildtype) while the phosphomimetic shows no significant difference in the number of cells with >10 foci over a mutant with all three QTs mutated (47±12% for T1171E vs 47±5% for LC8 binding null) (Fig. 7D,E). The similarity of the phosphodeficient mutant to the wildtype and the strong loss of foci in the phosphomimetic show that QT2 is not phosphorylated in response to ionizing radiation and argues against the importance of phosphorylation at this site in nuclear repair foci. Constructs with QT2 binding removed by AAA mutation (QT3 and QT13) fail to show a significant improvement in focus formation over the LC8 binding null mutant, while the mutant containing only QT2 does provide an enhancement. All of this together suggest a role for QT2 in regulating the enhancement from LC8 in focus formation of 53BP1, which we propose to be mediated through heterogeneous bridging of 53BP1 trimers.

## Discussion

53BP1 is a key player in the regulation of DNA damage repair, entering DNA repair foci upon induction of DNA damage and orchestrating selection of double-strand break repair pathway^8,11,45^. While the OD of 53BP1 is required for accumulation of 53BP1 in DNA repair foci, LC8 binding rescues some 53BP1 foci in the absence of the OD^10,12^. Mutation of LC8 binding sites results in DNA repair deficient 53BP1 and reduces the number of nuclear bodies formed upon induction of DNA damage, suggesting synergistic function between the 53BP1 OD and LC8^6^. Since LC8 functions as a dimerization hub, often regulating the dimerization of client proteins or contributing to higher-order oligomerization^28,29,46^, it was proposed that LC8 rescues 53BP1 foci by dimerizing 53BP1 in the absence of the OD. Here we provide the first evidence that 53BP1 is a trimer, not a dimer, that its interaction with LC8 forms a heterogeneous mixture containing bridged complexes, and that this bridging of two trimers by LC8 is what regulates 53BP1 activity.

### 53BP1 is a trimer and forms a dimer-of-trimers with LC8

While clients of LC8 are often dimers^29,31,32^, and occasionally tetramers^47,48^, there are no reports of trimeric clients. Interestingly we find that 53BP1 is a trimer and that LC8 can stimulate higher-order oligomerization by bridging two trimers of 53BP1 (Fig. 3B,D). A titration of 53BP1 LBD-OD analyzed by ^1^H-^15^N HSQC shows uniform peak loss at the three LC8 binding sites (Fig. 2F), suggesting that LC8 binds 53BP1 cooperatively and that there is no preference for any QT to be filled before other QTs. Results from AUC show heterogeneous complexes while SEC-MALS identifies multiple species with masses corresponding to LBD-OD bound to LC8 in trimer, bridged dimer-of-trimers, and intermediate species (Fig. 3B).

The difference in complexes observed by SEC-MALS and AUC indicate that while all QTs can generate trimer and bridged complexes, only QT2 is effective at forming bridged species stable enough to remain bound upon dilution during SEC (Fig. 4B,C). This is confirmed by QT mutants of LBD-OD, which showed SEC-MALS data consistent with a model in which either trimers or dimers-of-trimers are formed and all intact QTs are bound. Together, these data suggest a complex structure in which the 53BP1 OD trimers are bridged by LC8 and stabilized through interaction with QT2.

ITC supports a model in which QT2 is the primary motif that LC8 uses to bridge the 53BP1 trimers (Fig. 5). When titrating LC8 into LBD-OD mutants that retain a single LC8 site, QT2 is the one-site mutant that has the highest favorable enthalpy for LC8 binding (Fig. 5K); this is observed with both LBD and LBD-OD indicating it is a property of the motif. Additionally, all LBD-OD mutants with QT2 binding removed show isotherms consistent with two-step binding. These isotherm shapes are not seen in titrations of LBD alone. For example, QT13-OD would first form a stable trimer bound to 2 LC8 dimers, leaving one of the LBD chains completely unoccupied which upon further addition of LC8, will form a bridged trimer. When the titration is reversed (QT13-OD into LC8), there is no intermediate detected because the trimer complex is not stable in excess LC8 providing support for our model. (Fig. 5J). SBS fits of ITC data of one-site LBD-OD mutants show a small difference in affinity between bridging and intratrimer interactions only for QT2 (Fig. 5L) which explains the absence of an intermediate species in constructs containing QT2.

### LC8 binding forms heterogeneous 53BP1 complexes

LC8 is a hub protein involved in binding >100 client proteins at disordered regions^24,26^, facilitating their dimerization and regulating their physiological functions. While it is clearly established that binding to LC8 results in dimerization of the client^46^, the structural and physiological consequences of LC8 binding can be more complex. Rigidification of the disordered strand (Nup159)^30^, restriction of conformational ensemble (RavP)^32^, regulation of self-association (Swa)^28^, higher-order oligomerization (LCA5)^47^, and client regulation through cooperative binding (ASCIZ)^49,50^ are all mechanisms by which LC8 regulates its clients’ functions. Here we identify bridging of trimeric partner complexes of 53BP1 as a novel binding mode for LC8.

The outcome of the interaction between 53BP1 and LC8 is higher-order oligomerization of 53BP1, but there is an upper boundary to the size of the complex formed. Once 53BP1 forms the bridged dimer-of-trimers complex, there are no LBD chains available to bind (trimer 53BP1 leaves one free chain after LC8 binds) (Fig. 3D). A similar mechanism is seen in the interaction between LCA5 and LC8^47^. LCA5 has two LC8 binding sites and tetrameric coiled-coils, however the arrangement of these domains is unique for LC8 clients. The two coiled-coils are located on either side of the LC8 binding sites, and as a result each LC8 site is capable of bridging to a different LCA5 tetramer. Since there is always a free LC8 site to bridge to another LCA5 subcomplex, the higher-order oligomerization is unbounded, resulting in large assemblies of LCA5-LC8 that appear as beads-on-a-string by electron microscopy. While LCA5 and 53BP1 both form higher-order assemblies bridged by LC8 dimers, they form drastically different complexes, presumably due to their difference in native oligomeric state. The trimeric OD of 53BP1 results in bounded bridging with an upper limit of a dimer-of-trimers, while LCA5 exhibits unbounded bridging in complex with LC8.

Given that all LC8 sites are filled simultaneously and that QT2 appears to be sufficient for bridging, what is then the reason for multivalent sites in 53BP1-LC8 interactions? The binding enhancement from multivalent LC8 interactions is minimal in this case, contrary to what is observed with ASCIZ QT2-4^41,42^ and Nup159^30,31^ for example. WT LBD-OD has a K_d_ of 0.4 μM, while 2-site mutants have K_d_ between 0.4-0.7 μM. Either QT1 or QT3 could be absent and make almost no change to the overall affinity of the interaction (QT23-OD K_d_= 0.4 μM QT12=OD K_d_=0.5 μM). A model in which LC8 binding stabilizes bivalent contacts between 53BP1 and modified histones due to restriction of the conformational ensemble of 53BP1 has been proposed^6^ (Fig. 8A, 53BP1 dimer). Such a binding mode is seen in the dimeric LC8 binding client, RavP (Fig. 8A), where LC8 binding restrict its conformational ensemble^32^. While there certainly is some binding enhancement of LBD-OD compared to QT2-OD and could possibly be some ordering of the 53BP1 LBD upon binding LC8 multivalently, we propose a different reason for multivalency in 53BP1-LC8 interactions. While QT2 is both necessary and sufficient for bridging, deletion of part of the linker separating LBD and OD results in a primarily bridged complex (Fig. 6). We propose that the motif specificity of QT2 and the position of QT1 and QT3 contribute to tuning the affinity of the bridging interaction, optimizing the relative population of trimer and bridged complexes.

**Figure 8.**
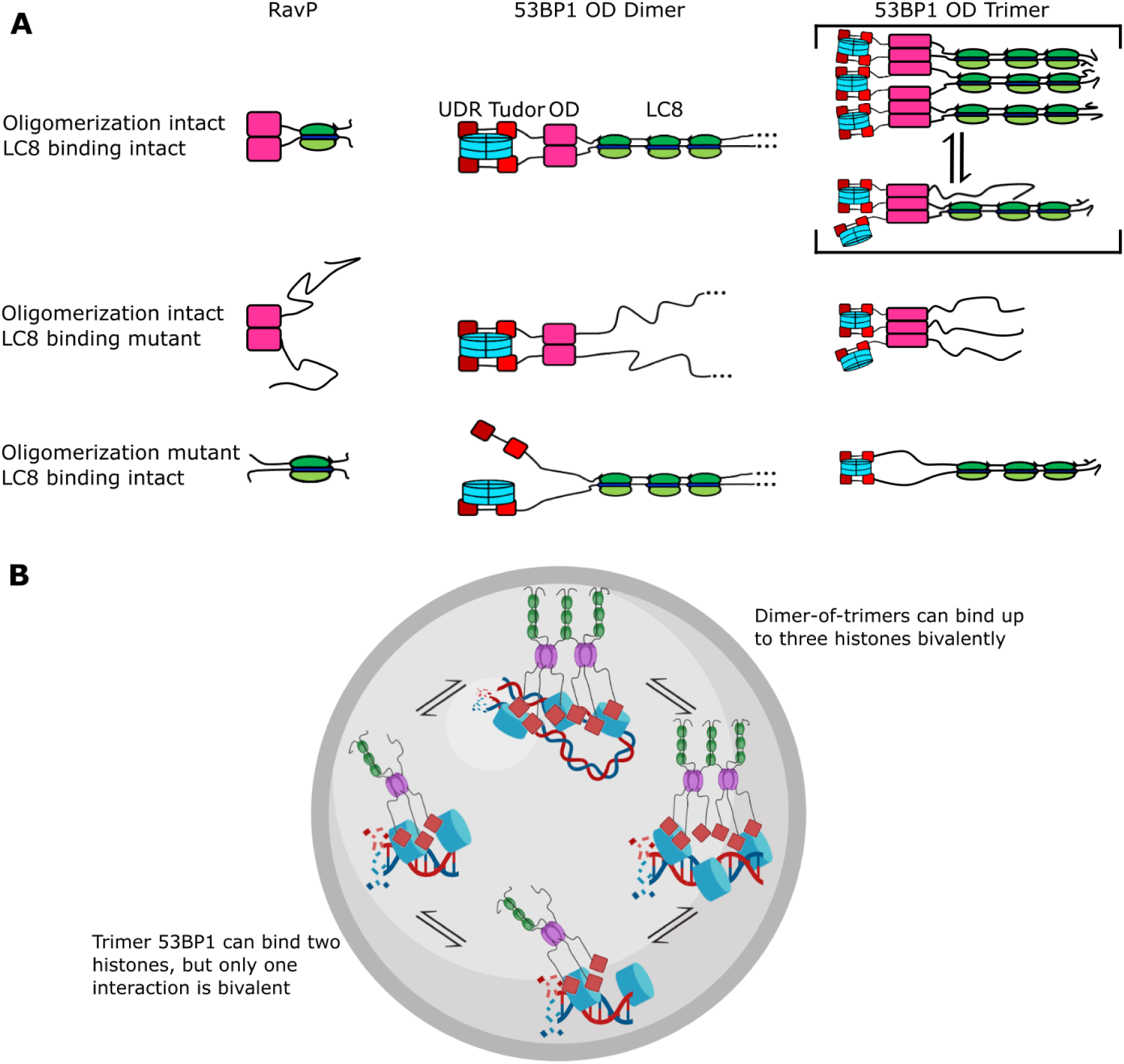
Mechanism of LC8 enhancement of 53BP1 focus formation. **A**, Cartoon showing roles of LC8 in interaction with RavP^32^ (left), published mechanisms for LC8 with 53BP1^6^ (center) and revised mechanism based on this work (right). In RavP, the N-termini are aligned with LC8, but when LC8 binding is removed the termini occupy a larger conformational ensemble. In the dimeric 53BP1 OD model, 53BP1 is a dimer as long as either the OD or LC8 binding is intact. This model does not fully capture the effect of OD and LC8 mutation of 53BP1 focus formation and cannot be correct because 53BP1 is a trimer not a dimer. An alternative model is the trimeric 53BP1 OD which results in higher-order oligomerization of 53BP1 with LC8. Mutation of the OD or LC8 binding would then be expected to produce the observed differences in reductions in focus formation in the ODm and LC8m. **B**, LC8-53BP1 binding and accumulation in DNA repair foci. 53BP1/LC8 complex is in exchange between trimeric and bridged species.

### Heterogeneity of oligomeric state regulates 53BP1 focus formation

Currently, 53BP1 is thought of as a dimer that can form higher-order oligomers^6,13,27^ and LC8 binding will stabilize the 53BP1 OD dimer, resulting in an increase in affinity for bivalent interactions with histones^6^ (Fig. 8A, 53BP1 dimer). Based on nuclear focus formation and chromatin-association of 53BP1 OD and LC8 mutants, loss of LC8 binding results in only a modest reduction in chromatin association compared to the wild type, while loss of the OD generates 53BP1 that is dimerized only by LC8 but has a significant reduction in chromatin association. Loss of both the OD and QT sites abolishes 53BP1 foci almost completely. While this model attributes the reduction of 53BP1 focus formation to the lack of alignment of the N-terminal disordered region of 53BP1 in LC8m, it does not accurately reflect the structural consequences of LC8 binding on higher-order oligomerization. Our model which includes trimer and bridged 53BP1 oligomers will account for the change in 53BP1 chromatin association because of the higher affinity for histones due to increased avidity (Fig. 8A, 53BP1 trimer). In this model, the LC8 binding deletion mutant is a trimer and the OD deletion mutant is a dimer, also explaining the modest and large reduction in 53BP1 foci for these constructs, respectively^6^. Therefore, our trimer and dimer-of-trimers model fully accounts for changes in 53BP1 function resulting from loss of LC8 binding and should provide a more accurate representation of 53BP1-LC8 interactions.

In summary, we show for the first time that 53BP1 OD is a homogeneous trimer. Binding to LC8 results in a heterogenous mixture of complexes with masses consistent with the formation of both trimer and bridged complexes of 53BP1. We speculate that heterogeneity in 53BP1-LC8 complexes may allow for improved recruitment of 53BP1 into nuclear repair foci, and the bridged complex stabilizes interactions with chromatin by allowing for more bivalent interactions (Fig. 8A). Both the trimer and bridged complexes can bind multiple histones, but the dimer-of-trimers could bind up to 3 histones (blue) bivalently, resulting in high avidity and retention in DNA repair foci (Fig. 8B). Indeed, mutants shown to be deficient in bridging 53BP1 trimers *in vitro* fail to elicit an improvement in 53BP1 focus formation while even the weakly bridging QT2 construct improved focus formation significantly compared to an LC8 binding null mutant, providing support for the physiological relevance of LC8 in bridging 53BP1 trimers (Fig. 6D,E). Our model (Fig. 8) explains previous data involving 53BP1-chromatin interactions and offers a new role for LC8 in the enhancement of 53BP1 foci.

## Methods

### Cloning and site directed mutagenesis

Studies were carried out using human 53BP1 (Uniprot: Q12888) and human LC8-2 (Uniprot: Q96FJ2). Plasmid containing 53BP1 1140-1290 was codon optimized for expression in Escherichia coli and cloned into a pET24d expression vector (GenScript, Piscataway, NJ, USA). All constructs contained an N-terminal hexahistidine tag and tobacco etch virus (TEV) protease cleavage site to facilitate removal of the his-tag. Cysteine residues in the sequence were mutated to serine to avoid disulfide induced oligomerization. All mutagenesis was performed using New England Biolabs’ (Ipswich, MA, USA) site-directed mutagenesis kit and custom primers. The wild-type 53BP1 LBD-OD contains all three LC8-binding sites, while the QT-OD variants contain only the indicated LC8-binding sites (i.e., QT1-OD contains intact LC8-binding site 1, while LC8-binding sites 2 and 3 are mutated to AAA to abolish binding). T1171 point mutants are LBD-OD variants containing only a single amino acid change, as indicated in the name of the mutant. Linker deletion mutants are LBD-OD variants with indicated truncation of residues from LBD-OD linker.

### Protein expression and purification

All constructs were transformed into E. coli Rosetta DE3 cells and expressed in rich autoinducing media at 37°C for 24 h or grown with Luria-Bertani and induced at 18°C for 16-20 h with 1 mM IPTG. For NMR experiments, cells were grown in MJ9 minimal media supplemented with ^13^C-glucose and/or ^15^NH_4_Cl. Cells were harvested and purified on Talon His-Tag Purification Resin (Takara Bio, Mountain View, CA, USA). The hexahistidine tag was removed by incubating overnight at 4°C with TEV protease. Cleavage with TEV protease leaves an N-terminal Gly-Ala-His preceding the peptide of interest. The cleaved samples were passed back over Talon resin to remove the cleaved tag and TEV protease. Samples were then purified on S200 Superdex size exclusion column using an AKTA-FPLC (GE Healthcare, Chicago, IL, USA) to a purity of >95%, as analyzed by SDS-PAGE. Proteins were stored at 4°C and either used within 1 week or flash frozen and stored at -80°C.

Protein concentration was quantified using absorbance at 280 and 205 nm since the 53BP1 LBD-OD sequence contains only two tyrosine residues using extinction coefficients of: at 205 nm LC8-2 374,720 M^-1^cm^-1^ and LBD-OD constructs (including mutants) 508,340 M^-1^cm^-1^; at 280 nm LC8-2 14440 M^-1^cm^-1^ and LBD-OD constructs (including mutants) 2,980 M^-1^cm^-1^.

### Circular dichroism

Spectra were recorded on a JASCO J-810 spectropolarimeter using a 1 mm cell. Protein samples were dialyzed overnight in 2 L of 20 mM sodium phosphate (pH 7.2) prior to data collection. Spectra were collected at room temperature at a protein concentration of 10-15 μM. Data shown are the average of 3 scans, and results are reported in mean residue molar ellipticity (deg*cm^2^/dmol).

For thermal denaturing assay, spectra were recorded at the specified temperatures (between 25°C and 80°C). A single sample was used for each assay without replacing the sample between points. Samples were allowed to thermally equilibrate for 10 minutes prior to data acquisition.

### Isothermal titration calorimetry

Thermodynamics of the 53BP1 LBD-OD:LC8 interaction were measured at 25°C using a VP-isothermal titration calorimetry (ITC) microcalorimeter (MicroCal, Northampton, MA, USA). The binding buffer contained 50 mM sodium phosphate and 150 mM sodium chloride (pH 7.2). LC8 at a concentration of 250–400 μM was titrated into an LBD-OD construct. Cell concentrations were between 10-35 μM for each LBD-OD mutant. For experiments where LBD-OD mutant was titrated into LC8, LBD-OD at a concentration of 300 μM was titrated into LC8 mutant at concentration on 10-15 μM. Peak areas were integrated, and the data were fit to a one-site binding model in Origin 7.0 (OriginLab Corporation, Northampton, MA, USA). From this, stoichiometry (N), dissociation constant (K_d_), change in enthalpy (ΔH), and entropy (ΔS) were determined. Reported data are the average of triplicate runs. Error reported is the standard deviation of the data acquired.

Each isotherm of the one-site mutants of LBD-OD was additionally analyzed using the Subsequent Binding Sites (SBS) models found within Origin for VP-ITC. We model that the hexamer binds three LC8 dimers, two of which bind with identical conditions (intratrimer; between two LBD chains within the same trimer) and one binds in a different condition (bridging; linking two 53BP1 trimer together). For SBS, the following constraints were applied: k1 = k2; h1 = h2; k1 > k3; h1 < h3. This fitting strategy was followed for every experimental replicate of each one-site mutant, and inverse-variance weighting (equations 1 & 2) was used to aggregate the measured values and their associated errors for each replicate. For a measurement yi and associated error σi, inverse-variance weighting for calculating an aggregated measurement (*y^*) and error (*σ^*) can be represented by the following equations:

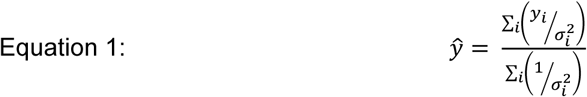

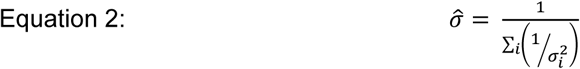

### Size-exclusion chromatography coupled to multi-angle light scattering

Average molar masses and association states of proteins were determined from SEC (AKTA FPLC; GE Healthcare) coupled to multi-angle light scattering (DAWN; Wyatt Technology) and refractive index (Optilab; Wyatt Technology) detectors. Size-exclusion chromatography was performed on S200 Superdex gel filtration column using an AKTA-FPLC (GE Healthcare, Chicago, IL, USA). Data for free 53BP1 LBD-OD or OD were collected by injecting protein at 1-10 mg/mL with the column pre-equilibrated with the ITC buffer described above. For binding experiments, 50 μM 53BP1 LBD-OD mutant was mixed with 200 μM LC8 and allowed to bind for 30 minutes at 4°C. Samples were run at a flow rate of 0.6 mL/min. Average molar masses and associated uncertainties were computed with the ASTRA software package, version 8 (Wyatt Technologies). Expected masses are as follows: 53BP1 LBD-OD 17.0 kDa, LC8 21.2 kDa, OD 13.2 kDa. Proteins concentration was measured using Optilab refractive index detector (Wyatt Technology, Santa Barbara, CA, USA).

### Sedimentation velocity analytical ultracentrifugation

Analytical ultracentrifugation (AUC) experiments were performed using a Beckman Coulter Optima XL-A ultracentrifuge equipped with absorbance optics (Brea, CA, USA). For sedimentation velocity AUC (SV-AUC) experiments, samples were loaded into epon 2-channel sectored cells with a 12 mm optical pathlength and run at 42,000 Rpm in a four-cell Beckman Coulter AN 60-Ti rotor at 20°C. Scans were performed continuously for a total of 300 scans per cell. Data were fit to a continuous c(S) distribution using the software SEDfit. Sedimentation coefficients are expressed in Svedbergs (S).

### NMR

NMR experiments were performed on a Bruker 800 MHz Avance III HD spectrometer (Bruker Biosciences, Billerica, MA, USA) equipped with a 5 mm TCI cryoprobe with Z-axis gradient (Bruker). NMR experiments were carried out at 10°C in 20 mM sodium phosphate, 50 mM sodium chloride, and 1 mM Sodium azide (pH 6.5), with 10% D2O, a protease inhibitor mixture (Roche Applied Science, Madison, WI, USA), and 2-2 dimethylsilapentane-5-sulfonic acid for chemical shift referencing. NMR data were processed using nmrPipe ^51^ and non-uniform sampling artifacts removed using SMILE ^52^. Backbone resonances were previously assigned in the LBD^23^ and have been deposited in the BMRB (BMRB entry 51475). Data analysis was performed on NMRViewJ^53^. Steady-state 1H-15N heteronuclear nuclear Overhauser effects ({1H}-15N NOE) experiments were collected with an 8 s saturation time. Error bars were calculated using:

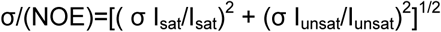

where I_sat_ and σ I_sat_ are the intensity of the peak and its baseline noise. R1 and R2 data were collected with heteronuclear single quantum coherence (HSQC)-based temperature compensated pulse sequences^54^. Time points were collected in triplicate for error estimation. For titration of LBD-OD with LC8, 1H-15N HSQC spectra of 100 μM ^15^N-labeled LBD-OD were collected with 0.25, 0.5, 1, 2, 3, and 4 molar equivalents of unlabeled LC8 titrated using the NMR conditions described above. Chemical shift indexing was performed using sequence-corrected shifts for amino acids in random coils^55–57^.

### Immunofluorescence staining

U2OS cells were transfected with 3 μg GFP-expression vectors using 10 μl polyethylenimine (1 mg/ml). Cells were irradiated using MultiRad (Precision XRay Irradiation) with indicated dose 24 hour after transfection followed by 1-hour rest. The coverslips were then fixed with 3% paraformaldehyde for 10 minutes and permeabilized using 0.5% Triton-X solution for 5 minutes. Samples were incubated with mouse monoclonal primary γH2AX antibody (EMD Millipore, 05-636) (1:1000) for 1 hour followed by Alexa Fluor 594 goat anti-mouse (1:500) and DAPI (1:5000) for 30 minutes. Coverslips were mounted using 0.02% anti-fade solution (0.02% in 90% glycerol). Samples were imaged using Zeiss LSM900 confocal microscope and analyzed using GraphPad Prism. Data represent 3-4 individual replicates.

## Data availability

All experimental data of the manuscript is available upon e-mail request from Elisar Barbar

## Acknowledgments

The authors would like to thank Austin Weeks for help generating reagents. We also thank Nikolaus Loening, Sanjay Ramprasad, and Sarah McGee for thoughtful conversation and editing. J.H. acknowledges funding from ARCS Oregon Chapter and the Dean’s Catalyst award. This work is funded by the National Institutes of Health (R01GM141733 to E.B.). We also acknowledge the support of the Oregon State University NMR Facility funded by the National Institutes of Health, HEI grant 1S10OD018518, and the M. J. Murdock Charitable Trust grant #2014162. J.W.L. is supported by grant from NIH (NIGMS: R35GM137798, NCI: R01CA244261), American Cancer Society (RSG-20-131-01-DMC and TLC-21-164-01-TLC) and University of Texas STARs award.

## References

1. Iwabuchi, K., Bartel, P. L., Li, B., Marraccino, R. & Fields, S. Two cellular proteins that bind to wild-type but not mutant p53. Proc. Natl. Acad. Sci. 91, 6098–6102 (1994).

2. Rappold, I., Iwabuchi, K., Date, T. & Chen, J. Tumor Suppressor P53 Binding Protein 1 (53bp1) Is Involved in DNA Damage–Signaling Pathways. J. Cell Biol. 153, 613–620 (2001).

3. Mirza-Aghazadeh-Attari, M. et al. 53BP1: A key player of DNA damage response with critical functions in cancer. DNA Repair 73, 110–119 (2019).

4. Gupta, A. et al. Role of 53BP1 in the regulation of DNA double-strand break repair pathway choice. Radiat. Res. 181, 1–8 (2014).

5. Ward, I. M., Minn, K., van Deursen, J. & Chen, J. p53 Binding protein 53BP1 is required for DNA damage responses and tumor suppression in mice. Mol. Cell. Biol. 23, 2556–2563 (2003).

6. Becker, J. R. et al. The ASCIZ-DYNLL1 axis promotes 53BP1-dependent non-homologous end joining and PARP inhibitor sensitivity. Nat. Commun. 9, 5406 (2018).

7. Bouwman, P. et al. 53BP1 loss rescues BRCA1 deficiency and is associated with triple-negative and BRCA-mutated breast cancers. Nat. Struct. Mol. Biol. 17, 688–695 (2010).

8. Bunting, S. F. et al. 53BP1 Inhibits Homologous Recombination in Brca1-Deficient Cells by Blocking Resection of DNA Breaks. Cell 141, 243–254 (2010).

9. Al-Ejeh, F. et al. Harnessing the complexity of DNA-damage response pathways to improve cancer treatment outcomes. Oncogene 29, 6085–6098 (2010).

10. West, K. L. et al. LC8/DYNLL1 is a 53BP1 effector and regulates checkpoint activation. Nucleic Acids Res. 47, 6236–6249 (2019).

11. Schultz, L. B., Chehab, N. H., Malikzay, A. & Halazonetis, T. D. P53 Binding Protein 1 (53bp1) Is an Early Participant in the Cellular Response to DNA Double-Strand Breaks. J. Cell Biol. 151, 1381–1390 (2000).

12. Kilic, S. et al. Phase separation of 53BP1 determines liquid-like behavior of DNA repair compartments. EMBO J. 38, (2019).

13. Zgheib, O., Pataky, K., Brugger, J. & Halazonetis, T. D. An Oligomerized 53BP1 Tudor Domain Suffices for Recognition of DNA Double-Strand Breaks. Mol. Cell. Biol. 29, 1050–1058 (2009).

14. Fradet-Turcotte, A. et al. 53BP1 is a reader of the DNA-damage-induced H2A Lys 15 ubiquitin mark. Nature 499, 50–54 (2013).

15. Kelliher, J. L. et al. Evolved histone tail regulates 53BP1 recruitment at damaged chromatin. Nat. Commun. 15, 4634 (2024).

16. Zgheib, O. et al. ATM signaling and 53BP1. Radiother. Oncol. 76, 119–122 (2005).

17. Jowsey, P. et al. Characterisation of the sites of DNA damage-induced 53BP1 phosphorylation catalysed by ATM and ATR. DNA Repair 6, 1536–1544 (2007).

18. Zimmermann, M., Lottersberger, F., Buonomo, S. B., Sfeir, A. & De Lange, T. 53BP1 Regulates DSB Repair Using Rif1 to Control 5′ End Resection. Science 339, 700–704 (2013).

19. Ghezraoui, H. et al. 53BP1 cooperation with the REV7–shieldin complex underpins DNA structure-specific NHEJ. Nature 560, 122–127 (2018).

20. Noordermeer, S. M. et al. The shieldin complex mediates 53BP1-dependent DNA repair. Nature 560, 117–121 (2018).

21. Wu, J., Prindle, M. J., Dressler, G. R. & Yu, X. PTIP Regulates 53BP1 and SMC1 at the DNA Damage Sites. J. Biol. Chem. 284, 18078–18084 (2009).

22. Lo, K. W.-H. et al. The 8-kDa Dynein Light Chain Binds to p53-binding Protein 1 and Mediates DNA Damage-induced p53 Nuclear Accumulation *. J. Biol. Chem. 280, 8172–8179 (2005).

23. Howe, J., Weeks, A., Reardon, P. & Barbar, E. Multivalent binding of the hub protein LC8 at a newly discovered site in 53BP1. Biophys. J. 121, 4433–4442 (2022).

24. Jespersen, N. et al. Systematic identification of recognition motifs for the hub protein LC8. Life Sci. Alliance 2, e201900366 (2019).

25. Benison, G., Karplus, P. A. & Barbar, E. Structure and dynamics of LC8 complexes with KXTQT-motif peptides: swallow and dynein intermediate chain compete for a common site. J. Mol. Biol. 371, 457–468 (2007).

26. Barbar, E. & Nyarko, A. Polybivalency and disordered proteins in ordering macromolecular assemblies. Semin. Cell Dev. Biol. 37, 20–25 (2015).

27. Lou, J., Priest, D. G., Solano, A., Kerjouan, A. & Hinde, E. Spatiotemporal dynamics of 53BP1 dimer recruitment to a DNA double strand break. Nat. Commun. 11, 5776 (2020).

28. I. Kidane, A., et al. Structural Features of LC8-Induced Self-Association of Swallow. Biochemistry 52, 6011–6020 (2013).

29. Barbar, E. & Nyarko, A. NMR CHARACTERIZATION OF SELF-ASSOCIATION DOMAINS PROMOTED BY INTERACTIONS WITH LC8 HUB PROTEIN. Comput. Struct. Biotechnol. J. 9, e201402003 (2014).

30. Nyarko, A., Song, Y., Nováček, J., Žídek, L. & Barbar, E. Multiple Recognition Motifs in Nucleoporin Nup159 Provide a Stable and Rigid Nup159-Dyn2 Assembly. J. Biol. Chem. 288, 2614–2622 (2013).

31. Gaik, M. et al. Structural basis for assembly and function of the Nup82 complex in the nuclear pore scaffold. J. Cell Biol. 208, 283–297 (2015).

32. Jespersen, N. E. et al. The LC8-RavP ensemble Structure Evinces A Role for LC8 in Regulating Lyssavirus Polymerase Functionality. J. Mol. Biol. 431, 4959–4977 (2019).

33. Erdős, G., Pajkos, M. & Dosztányi, Z. IUPred3: prediction of protein disorder enhanced with unambiguous experimental annotation and visualization of evolutionary conservation. Nucleic Acids Res. 49, W297–W303 (2021).

34. Benison, G., Karplus, P. A. & Barbar, E. The interplay of ligand binding and quaternary structure in the diverse interactions of dynein light chain LC8. J. Mol. Biol. 384, 954–966 (2008).

35. Kollmar, M. & Simm, D. Benchmark analysis of coiled-coil prediction tools. 486501065 Bytes figshare 10.6084/M9.FIGSHARE.9994706 (2019).

36. Delorenzi, M. & Speed, T. An HMM model for coiled-coil domains and a comparison with PSSM-based predictions. Bioinformatics 18, 617–625 (2002).

37. Micsonai, A. et al. Accurate secondary structure prediction and fold recognition for circular dichroism spectroscopy. Proc. Natl. Acad. Sci. 112, (2015).

38. Jumper, J. et al. Highly accurate protein structure prediction with AlphaFold. Nature 596, 583–589 (2021).

39. McGuffin, L. J., Bryson, K. & Jones, D. T. The PSIPRED protein structure prediction server. Bioinformatics 16, 404–405 (2000).

40. Drozdetskiy, A., Cole, C., Procter, J. & Barton, G. J. JPred4: a protein secondary structure prediction server. Nucleic Acids Res. 43, W389–W394 (2015).

41. Walker, D. R. et al. Linker Length Drives Heterogeneity of Multivalent Complexes of Hub Protein LC8 and Transcription Factor ASCIZ. Biomolecules 13, 404 (2023).

42. Reardon, P. N. et al. The dynein light chain 8 (LC8) binds predominantly “in-register” to a multivalent intrinsically disordered partner. J. Biol. Chem. 295, 4912–4922 (2020).

43. Erdős, G. et al. Novel linear motif filtering protocol reveals the role of the LC8 dynein light chain in the Hippo pathway. PLOS Comput. Biol. 13, e1005885 (2017).

44. Clark, S., Nyarko, A., Löhr, F., Karplus, P. A. & Barbar, E. The Anchored Flexibility Model in LC8 Motif Recognition: Insights from the Chica Complex. Biochemistry 55, 199–209 (2016).

45. Lottersberger, F., Bothmer, A., Robbiani, D. F., Nussenzweig, M. C. & de Lange, T. Role of 53BP1 oligomerization in regulating double-strand break repair. Proc. Natl. Acad. Sci. 110, 2146 (2013).

46. Barbar, E. Dynein Light Chain LC8 Is a Dimerization Hub Essential in Diverse Protein Networks. Biochemistry 47, 503–508 (2008).

47. Szaniszló, T. et al. The interaction between LC8 and LCA5 reveals a novel oligomerization function of LC8 in the ciliary-centrosome system. Sci. Rep. 12, 15623 (2022).

48. Rodriguez Galvan, J., et al. Human Parainfluenza Virus 3 Phosphoprotein Is a Tetramer and Shares Structural and Interaction Features with Ebola Phosphoprotein VP35. Biomolecules 11, 1603 (2021).

49. Rapali, P. et al. LC8 dynein light chain (DYNLL1) binds to the C-terminal domain of ATM-interacting protein (ATMIN/ASCIZ) and regulates its subcellular localization. Biochem. Biophys. Res. Commun. 414, 493–498 (2011).

50. Clark, S. et al. Multivalency regulates activity in an intrinsically disordered transcription factor. eLife 7, e36258 (2018).

51. Delaglio, F. et al. NMRPipe: A multidimensional spectral processing system based on UNIX pipes. J. Biomol. NMR 6, (1995).

52. Ying, J., Delaglio, F., Torchia, D. A. & Bax, A. Sparse multidimensional iterative lineshape-enhanced (SMILE) reconstruction of both non-uniformly sampled and conventional NMR data. J. Biomol. NMR 68, 101–118 (2017).

53. Johnson, B. A. Using NMRView to Visualize and Analyze the NMR Spectra of Macromolecules. in Protein NMR Techniques vol. 278 313–352 (Humana Press, New Jersey, 2004).

54. Farrow, N. A. et al. Backbone Dynamics of a Free and a Phosphopeptide-Complexed Src Homology 2 Domain Studied by 15N NMR Relaxation. Biochemistry 33, 5984–6003 (1994).

55. Kjaergaard, M. & Poulsen, F. M. Sequence correction of random coil chemical shifts: correlation between neighbor correction factors and changes in the Ramachandran distribution. J. Biomol. NMR 50, 157–165 (2011).

56. Schwarzinger, S. et al. Sequence-Dependent Correction of Random Coil NMR Chemical Shifts. J. Am. Chem. Soc. 123, 2970–2978 (2001).

57. Kjaergaard, M., Brander, S. & Poulsen, F. M. Random coil chemical shift for intrinsically disordered proteins: effects of temperature and pH. J. Biomol. NMR 49, 139–149 (2011).

